# The putative Gαq-inhibiting pepducin P4Pal_10_ distinctly modulates function of the Gαi-coupled receptors FPR2 and FFA2R in neutrophils

**DOI:** 10.1101/745596

**Authors:** André Holdfeldt, Simon Lind, Claes Dahlgren, Huamei Forsman

**Author notes:** Corresponding author: André Holdfeldt, Department of Rheumatology and Inflammation Research, University of Gothenburg, Gothenburg, Sweden, (AH).

## Abstract

A novel mechanism of action was described, when a protease-activated receptor 4 (P4Pal_10_) derived lipopeptide (pepducin) was shown to inhibit signaling downstream of several unrelated Gαq-coupled receptors. We show that this putative Gαq-inhibiting pepducin lacks inhibitory effects on signaling downstream of the Gαq-coupled receptors for ATP (P2Y_2_R) and PAF (PAFR) expressed in human neutrophils. P4Pal_10_ inhibited however, signaling in neutrophils activated with agonists for the Gαi-coupled formyl peptide receptor 2 (FPR2) but not the closely related FPR1. In addition, the P4Pal_10_ pepducin was turned into an activating agonist in the presence of an allosteric modulator selective for free fatty acid receptor 2 (FFA2R). The results presented thus reveal Gαq-independent effects of P4Pal_10_ in modulating FPR2- and FFA2R-mediated neutrophil activation.

## Introduction

Receptors belonging to the G-protein-coupled receptor (GPCR) family constitute a class of recognition molecules involved in a broad range of physiological processes as well as pathological conditions (1). The GPCRs have in common that they are polypeptides that span a cell membrane seven times, and the transmembrane-spanning α-helices connect the N-terminal tail and three loops expressed on the cell surface, with three intracellular loops and a C-terminal tail facing the cytosolic compartment (2). To mediate a cellular response following recognition of an extracellular ligand that binds to its membrane exposed binding cavity/domain, the cytosolic domains are structurally and functionally modified to allow and mediate an intracellular coupling to a so-called heterotrimeric G-protein containing an α- and a β/γ subunit (3). There are four classes of α-subunits, designated as Gαs, Gαq, Gαi/o and Gα12/13, which initiate distinct as well as overlapping signaling cascades (3). The Gα subunit part possesses an intrinsic GTPase activity which keeps signaling of a receptor at a very low or zero level in the absence of an agonist (4). The conformational changes in the receptor, initiated by the binding of an agonist, promotes an exchange of GDP for GTP, which leads to dissociation and activation of the Gα subunit and a start of the G-protein dependent signaling cascade (3). The large structural similarities between G-proteins makes it hard to determine the precise identity of a signaling partner, and two bacterial toxins (pertussis toxin from *Bordetella pertussis* and cholera toxin from *Vibro cholera*) have for long been the only tools available to investigate receptor coupling to Gαi (sensitive to pertussis toxin) and Gαs (sensitive to cholera toxin), respectively (5, 6). New and more efficient inhibitors of specific G-protein subunits have been eagerly awaited and small molecule inhibitors of Gαq are now available (7, 8). These inhibitors (the depsipeptide YM-254890 and the analog FR900359) selectively target Gαq signaling through an inhibition of the GDP/GTP exchange, and signaling is thus selectively terminated downstream of the Gαq-protein.

Pepducins, lipidated peptides with an amino acid sequence identical to one of the intracellular parts of the GPCR to be targeted, represent a novel group of modulators (9, 10). According to the proposed model for interaction, the lipid moiety of a pepducin facilitates its transport over the cell membrane and the peptide part interacts with an allosteric receptor site exposed to the cytosol and binding of the pepducin either activates or inhibits signaling by the cognate receptor (9). According to the concept, pepducins should selectively target receptors that harbor a sequence in one of their cytosolic parts that is identical to that in the pepducin (10). Our earlier studies with GPCRs expressed by human neutrophils including the formyl peptide receptors (FPR1 and FPR2) and the ATP receptor (P2Y_2_R) have raised questions about the precise mechanism for how pepducins specifically activate/inhibit neutrophil GPCRs, and it is clear from these studies that the receptor selectivity is not always dictated solely by amino acid similarities between the receptor and the activating/inhibiting pepducin (1). In agreement with this, it was recently shown that the pepducin P4Pal_10_, a lipopeptide with an amino acids sequence identical to a part of the third intracellular loop of the protease-activated receptor 4 (PAR4), inhibits signaling not only by this receptor but also by an array of other GPCRs that signal through a G-protein containing a Gαq subunit (11). The P4Pal_10_ pepducin lacked effect on signaling by Gαs-coupled as well as Gαi-coupled receptors, and even though it had no direct effect on Gαq function, it was proposed to be a selective inhibitor for a group of receptors that recruit and signal through the Gαq-class of G-proteins (11).

The Gαq preference for P4Pal_10_ among different GPCRs in human neutrophils is not known, and we have now determined the effects of the presumed Gαq-inhibiting pepducin P4Pal_10_ on signaling induced by both Gαq- (PAFR and P2Y2R), and Gαi-coupled GPCRs (FFA2R, FPR1, and FPR2) expressed in neutrophils. We show that no inhibition was induced by P4Pal_10_ on the signals generated by Gαq-coupled receptors, whereas the responses triggered by the Gαi-coupled FPR2 and FFA2R was selectively inhibited and amplified, respectively.

## Material and Methods

### Ethics Statement

In this study, conducted at the Sahlgrenska Academy in Sweden, buffy coats obtained from the blood bank at Sahlgrenska University Hospital, Gothenburg, Sweden were used to isolate neutrophils. According to the Swedish legislation section code 4§ 3p SFS 2003:460, no ethical approval was needed since the buffy coats were provided anonymously.

### Chemicals

Dextran and Ficoll-Paque were obtained from GE-Healthcare Bio-Science (Uppsala, Sweden). The tripeptide fMLF, and bovine serum albumin, were purchased from Millipore Sigma (Burlington, MA, USA). Horseradish peroxidase (HRP) was obtained from Boehringer Mannheim (Germany). The hexapeptide WKYMVM, isoluminol, TNFα, propionate, dimethyl sulfoxide (DMSO) and ATP were obtained from Sigma (St. Louis, MO, USA). Cyclosporin H was kindly provided by Novartis Pharma (Basel, Switzerland) and PAF was from Avanti Polar Lipids Inc. (Alabama, USA). The pepducin F2Pal_10_ (12) and the palmitoylated ten amino acids long pepducin P4Pal_10_ (Pal-SGRRYGHALR) were synthesized by CASLO Laboratory (Lyngby, Denmark). The FPR2 peptidomimetic agonist Compound 14 (13) and the FPR2 peptidomimetic antagonist CN6 (14) were kind gifts from Henrik Franzyk (Copenhagen, Denmark). MCT-ND4 was a kind gift from Hidehito Mukai (Nagahama Institute of Bio-Science, Japan). PSMα2 was from the American Peptide Company (Sunnyvale, CA). YM-254890 was purchased from Wako Chemicals (Neuss, Germany). CATPB (FFA2R antagonist) was from Tocris Bioscience (Bristol, England). Fura-2 was from Thermo Fisher Scientific (Waltham, MA, USA). Cy5-WKYMVM was purchased from Phoenix Pharmaceuticals (Burlingame, CA).

All reagents were dissolved in DMSO and then further diluted in Krebs-Ringer phosphate buffer (KRG, pH 7.3; 120 mM NaCl, 5 mM KCl, 1.7 mM KH_2_PO_4_, 8.3 mM NaH_2_PO_4_ and 10 mM glucose) supplemented with Ca^2+^ (1 mM) and Mg^2+^ (1.5 mM).

### Isolation of human neutrophils and monocytes

Neutrophils from healthy donors were isolated from peripheral blood or buffy coats using dextran sedimentation and Ficoll-Paque gradient centrifugation as described (15). After hypotonic lysis of the remaining erythrocytes, neutrophils were washed, suspended in KRG, and stored on ice until use. Monocytes were isolated with dextran sedimentation and Ficoll-Paque, follow by negative selection with magnetic beads (16).

### Neutrophil NADPH-oxidase activity

The NADPH-oxidase activity was determined using isoluminol-enhanced chemiluminescence (CL) (17, 18) and measured in a six-channel Biolumat LB 9505 (Berthold Co., Wildbad, Germany). Disposable polypropylene tubes containing a 900 µl reaction mixture of 10^5^ neutrophils in KRG, isoluminol (2 × 10^−5^ M) and HRP (4 Units/mL), were equilibrated at 37°C for 5 minutes in the absence or presence of inhibitors before addition of 100 µL of stimulus.

### Calcium mobilization

Neutrophils were loaded with Fura-2-AM (5 µM) for 30 minutes in darkness at room temperature (RT) before washing and suspension in KRG. Measurements of intracellular calcium were carried out at 37°C in a PerkinElmer fluorescence spectrophotometer (LC50, Perkin Elmer, USA), with excitation wavelengths of 340 and 380 nm, an emission wavelength of 509 nm, and slit widths of 5 and 10 nm. The transient rise in intracellular calcium is presented as the ratio of fluorescence intensities (340 nm: 380 nm) detected.

### Cy5-WKYMVM competitive receptor binding assay

The pepducin P4Pal_10_ (2 µM) was added to neutrophils (1 × 10^6^ cells/ml) in KRG supplemented with Ca^2+^ and incubated on ice for 10 min, after which the fluorescently labeled FPR2 agonist Cy5-WKYMVM was added (1 nM), and incubation was continued for 1 h. Samples with Cy5-WKYMVM in the presence (non-specific binding) or absence (total binding) of WKYMVM (100 nM) were used as controls. The samples were analyzed by flow cytometry using an Accuri flow cytometer.

### Data analysis

Data analysis was performed using GraphPad Prism 8.1.0 (Graphpad Software, San Diego, CA, USA). Curve fitting was performed by non-linear regression using the sigmoidal dose-response equation (variable-slope). Student’s *t*-test was used for statistical analysis, \**p* < 0.05, \*\**p* < 0.01.

## Result

### The transient increase in the cytosolic concentration of free Ca^2+^ mediated by Gαq-coupled PAFR and P2Y_2_R, is not inhibited by the P4Pal_10_ pepducin

One of the very early events in neutrophils following activation by G-protein coupled receptor (GPCR) selective agonists, is the induction of the PLC-PIP_2_-IP_3_ intracellular signaling pathway leading to an increase in the intracellular/cytosolic concentration of free Ca^2+^ ([Ca^2+^]_i_) (19). The Gαq-coupled GPCRs for the lipid chemoattractant platelet activating factor (PAFRs) and for ATP (P2Y_2_Rs) are abundantly expressed by neutrophils. In accordance with the general signaling scheme, stimulation of these GPCRs with their respective receptor specific agonist induced a transient rise in [Ca^2+^]_i_ (Fig. 1A, B). The cyclic desipeptide YM-254890, an inhibitor that selectively inhibit signaling by GPCRs coupled to G-protein containing the Gαq-subunit, abolished the PAF-mediated as well as the ATP-induced rise in [Ca^2+^]_i_ (Fig. 1A, B), confirming that both P2Y_2_R and PAFR couple to Gαq (20, 21).

**Figure 1.**
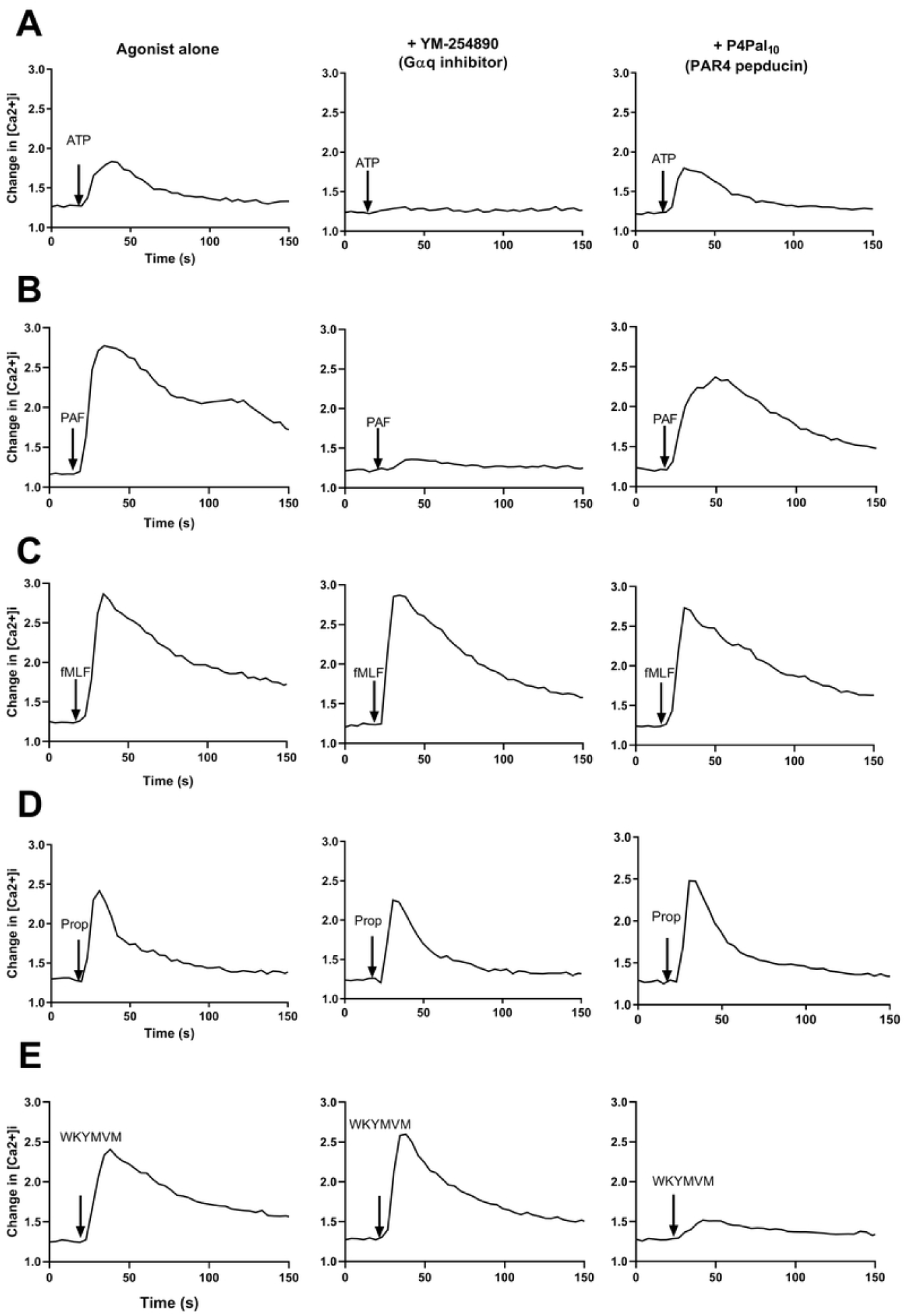
The P4Pal_10_ pepducin inhibits an increase of intracellular Ca^2+^ downstream of the FPR2 but not the Gαq-coupled PAFR and P2Y_2_-R in neutrophils. Neutrophils labeled with Fura2 were incubated (10 min, 37°C) without (agonist alone, left panels) or with YM-254890 (200 nM, middle panels) or P4Pal_10_ (500 nM, right panels). The transient rise in cytosolic Ca^2+^ was recorded upon stimulation with **A)** the P2Y2R agonist ATP (5 µM); **B)** the PAFR agonist PAF (0.5 nM); **C)** the FPR1 agonist fMLF (5 nM); **D)** the FFA2R agonist propionate (1.25 mM) **E)** the FPR2 agonist WKYMVM (5 nM). Arrows indicate agonist addition and data are presented as the ratio of fluorescence intensities (340 nm: 380 nm). A representative experiment is shown, n > 3. Abscissa: time in seconds (s), Ordinate: fluorescence (arbitrary units AU).

The PAR4-derived pepducin P4Pal_10_ has been shown to be a promiscuous inhibitor of GPCR-signaling that affects signaling by different Gαq-coupled receptors (11). Mechanistically, the P4Pal_10_ pepducin effect is indirect in that there is no direct effect on the function of the Gαq-protein; instead, the signaling capacity of sensitive receptors is modulated by the pepducin (11). As illustrated by the complete inhibition by YM-254890, the PAFRs and P2Y_2_Rs exclusively couple to Gαq, but despite this, no inhibition of the PAF- or ATP-induced transient rise in [Ca^2+^]_i_ was mediated by the P4Pal_10_ pepducin determined at concentrations up to 500 nM (Fig. 1A and B).

### The transient increase in [Ca^2+^]_i_ triggered by WKYMVM, an agonist specifically recognized by the Gαi-coupled FPR2, is selectively inhibited by the P4Pal_10_ pepducin

In addition to P2Y_2_R and PAFR, neutrophils express also pattern recognition GPCRs including FPR1/2 and the FFA2R that sense formylated peptides and free fatty acids such as propionate produced in the gut during microbial fermentation of fiber diets, respectively (22). The PLC-PIP_2_-IP_3_ intracellular signaling pathways leading to an increase of the cytosolic concentration of [Ca^2+^]_i_ was triggered also by the FPR1 agonists fMLF, the FPR2 agonist WKYMVM as well as by the FFA2R agonist propionate (Fig 1C-E). In agreement with the known Gαi preference (23), the Gαq selective inhibitor YM-254890 was without effect on the responses triggered by the agonist occupied FPRs and FFA2R (Fig 1 C-E). Similar to YM-254890, the P4Pal_10_ pepducin was without effect on the fMLF/FPR1-as well as on the propionate/FFA2R-induced transient rise in [Ca^2+^]_i_ (Fig 1C, D). The WKYMVM response was, however, substantially reduced in the presence of P4Pal_10_ (Fig 1E). Taken together, our data show that P4Pal_10_ has no effect on signaling by the Gαq-coupled PAFR and P2Y2R, but it selectively inhibits signaling by Gαi-coupled FPR2. The inhibitory effect of P4Pal_10_ is apparently not due to an inhibition of Gαi, a conclusion drawn from the fact that the pepducin had no inhibitory effect on signaling by the closely related FPR1 or by FFA2R in neutrophils.

### The FPR2-mediated activation of the superoxide (O_2_^−^) generating neutrophil NADPH-oxidase is selectively inhibited by the P4Pal_10_ pepducin

We next examined the effect of P4Pal_10_ on activation of the O_2_^−^ generating NADPH-oxidase, an electron transporting enzyme system in neutrophils that is assembled/activated by many agonist-occupied GPCRs (24, 25). Accordingly, agonists for the Gαi-coupled FPRs and for the Gαq-coupled PAFR are potent activators of the NADPH-oxidase, while ATP or propionate are unable to directly trigger an activate neutrophil NADPH-oxidase (26, 27). The neutrophil response induced by FPR- or PAFR-agonists has a rapid onset, reach a peak after around 1 min and is terminated in around 5 min (Fig. 2A-C). In agreement with the inability of the Gαq inhibitor YM-254890 to reduce the rise in [Ca^2+^]_i_ when triggered by FPR agonists (Fig 1C, E), no reduction in O_2_^−^ production was obtained in the presence of YM-254890 (Fig. 2A and B). As expected, the PAF-induced activation of NADPH-oxidase was largely inhibited by YM-254890 (Fig 2C). The P4Pal_10_ pepducin did not reduce, but rather significantly increased PAF-induced production of O_2_^−^ (Fig 2C and D). In line with the data obtained in Ca^2+^ assay system, P4Pal_10_ selectively inhibited FPR2-triggered activation of the NADPH-oxidase, as illustrated by the fact that the WKYMVM-induced production of O_2_^−^ was totally inhibited by the pepducin while the response induced by the the agonist for the closely related FPR1 was unaffected (Fig 2A and B). The effects of the FPR2 specific antagonist CN6 (14) was included for comparison (Fig 2A, B).

**Figure 2.**
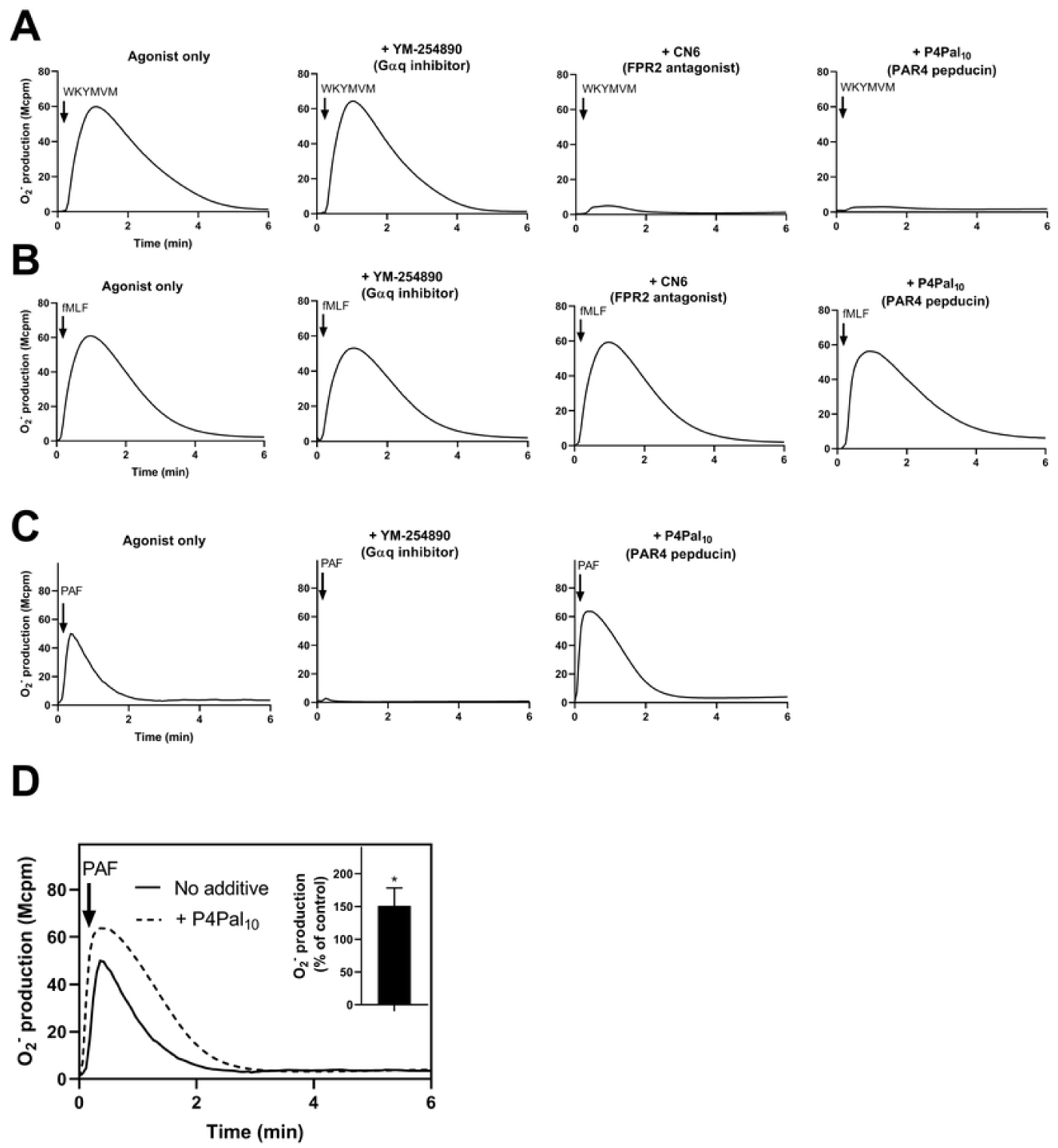
The P4Pal_10_ pepducin inhibits FPR2-but primes PAFR-mediated O_2_^−^ release from neutrophils. Neutrophils (10^5^) were incubated (5 min, 37°C) without (agonist only, left panels in A-C) or with YM-254890 (200 nM, Gαq inhibitor), CN6 (100 nM, FPR2 antagonist) or P4Pal_10_ (2 µM, PAR4 pepducin) before stimulation with agonist **A)** WKYMVM (100 nM); **B)** fMLF (100 nM); **C)** PAF (100 nM). The O_2_^−^ production was measured by isoluminol-amplified chemiluminescence over time. A representative experiment is shown, n > 3. Abscissa: time (Min), Ordinate: O_2_^−^ production (arbitrary units, Mcpm). **D).** A representative figure of P4Pal_10_-induced priming of the PAF response. Neutrophils (10^5^) were incubated (5 min, 37°C) without (solid line) or with P4Pal_10_ (2 µM, dotted line) followed by stimulation with PAF (100 nM). The O_2_^−^ production was measured by chemiluminescence over time. A representative experiment is shown, n = 3. Inset, quantification of the priming effect induced by the P4Pal_10_ pepducin on the PAF response. The data are expressed as % of control response (peak O_2_^−^ response induced by PAF in the absence of P4Pal_10_), (means ± sd; n = 3). \**P* < 0.05.

### The P4Pal_10_ pepducin inhibits O_2_^−^ production triggered also by other FPR2 agonists

The P4Pal_10_ pepducin abolished WKYMVM-induced O_2_^−^ production, and the inhibition was P4Pal_10_ concentration dependent with an IC_50_-value of ∼ 0.7 µM (Fig 3A). To further investigate whether the inhibitory effect of P4Pal_10_ is restricted to the agonist WKYMVM or achieved on the level of FPR2, we used several other FPR2 selective agonists including the *Staphylococcus aureus*-derived PSMα2 peptide (28), the mitochondrial-derived MCT-ND4 peptide (29), the lipidated peptidomimetic agonist Compound 14 (13), and the activating FPR2 specific pepducin F2Pal_10_ (12, 30). All these FPR2 agonists activate neutrophils and P4Pal_10_ inhibited the response (Fig 3B). These data clearly show that the P4Pal_10_ pepducin inhibits FPR2-mediated neutrophil activation, and the inhibition is at the level of receptor rather than on the specific ligand used.

**Figure 3.**
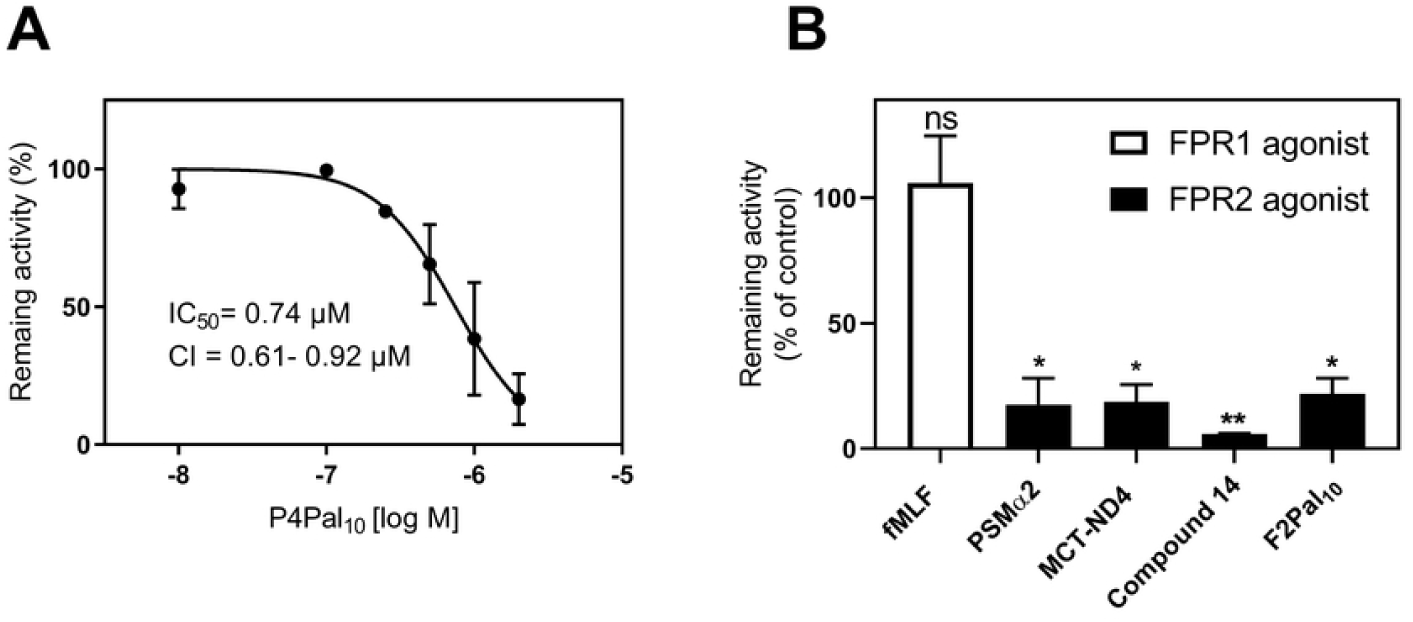
The P4Pal_10_ pepducin inhibits neutrophil O_2_^−^ release triggered by an array of FPR2 agonists. **A).** P4Pal_10_ dose-dependent inhibition of the neutrophil NADPH-oxidase activity induced by WKYMVM (100 nM). The results are presented as normalized WKYMVM responses, determined as the amount of O_2_^−^ produced (peak values) in the presence of increasing concentrations of P4Pal_10_, compared with production in the absence of P4Pal_10_. The IC_50_ value was calculated from three independent experiments. Abscissa, log concentration of P4Pal_10_; ordinate, peak value of O_2_^−^ production expressed as the percentage of the peak response obtained without P4Pal_10_ (means ± sd; n = 3). **B).** Effect of the P4Pal_10_ pepducin (2 µM) on neutrophil O_2_^−^ production induced by fMLF (100 nM), PSMα2 (50 nM), MCT-ND4 (5 nM), compound 14 (100 nM), and F2Pal_10_ (500 nM). The results are presented as remaining activity in the presence of the P4Pal_10_ pepducin, expressed as a percentage of the control activity (peak value) induced in the absence of P4Pal_10_ (means ± sd; n = 3). \**p* < 0.05, **< 0.01.

### The P4Pal_10_ pepducin blocked FPR2-mediated activation also in monocytes but lacked effect on binding of WKYMVM to FPR2

In addition to neutrophils that abundantly express FPR1 and FPR2, also monocytes express the two members of the FPRs in addition to the third family member FPR3 (23). To determine whether the P4Pal_10_ pepducin selectively targets FPR2-but not FPR1-signaling also in monocytes, we purified human blood-derived monocytes using CD14 negative selection magnetic beads and the monocyte purity was confirmed using CD14 antibody (Fig 4 A). Very similar to the results obtained from neutrophils, the P4Pal_10_ pepducin preferentially inhibited also the FPR2 but not FPR1-mediated O_2_^−^ production when monocytes were used (Fig 4B, C). Despite the clear inhibitory effect of the P4Pal_10_ pepducin on FPR2-mediated responses in both neutrophils and monocytes, our experiments in which P4Pal_10_ competes with a Cy5-labeled WKYMVM in binding to cell membrane exposed FPR2, show that the pepducin does not affect agonist binding (Fig 4D). Binding of Cy5-WKYMVM to FPR2 was, however, inhibited by non-labeled WKYMVM in large excess (Figure 4D).

**Figure 4.**
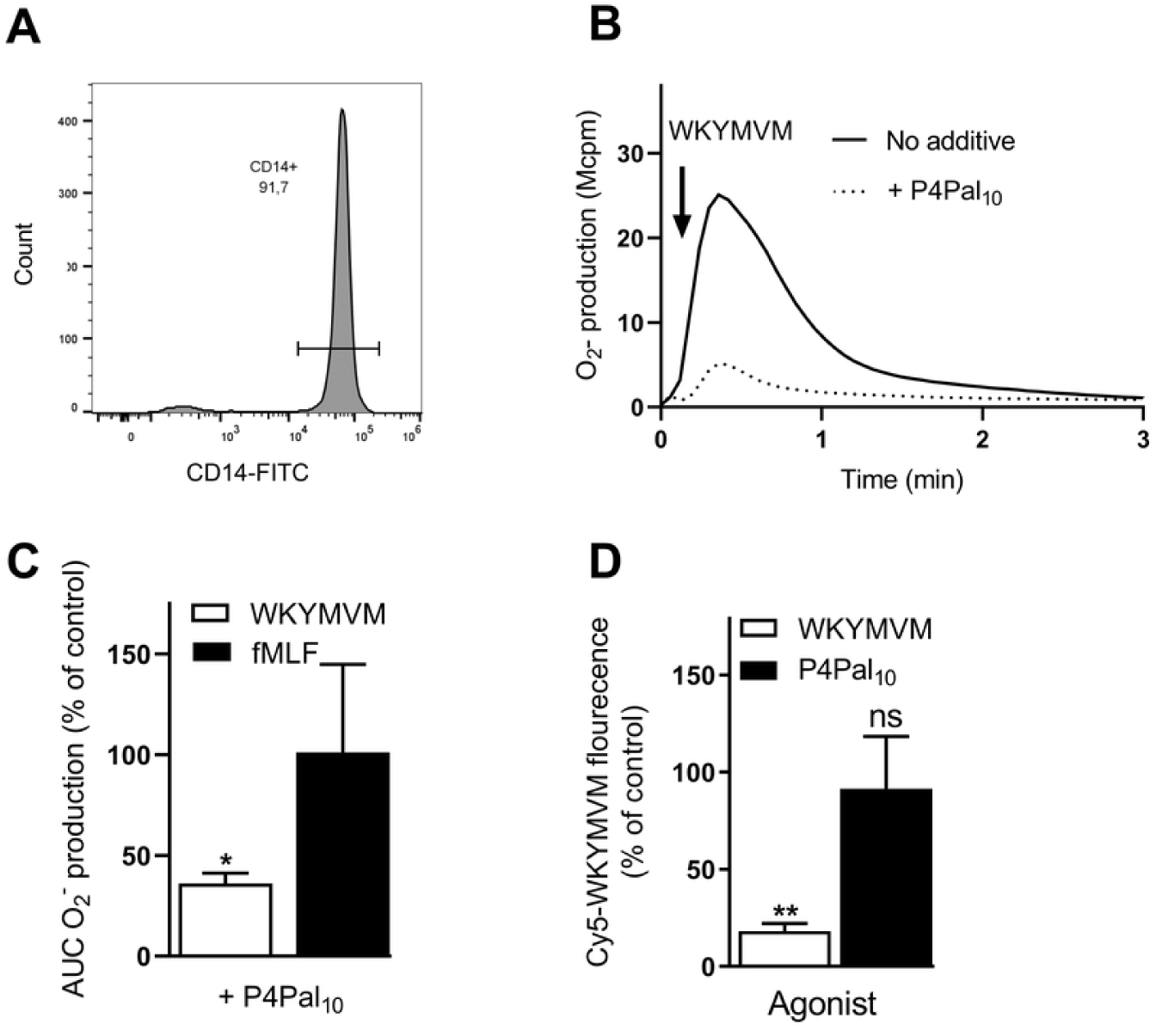
The P4Pal_10_ pepducin does not compete with WKYMVM for binding and is also an FPR2 antagonist in monocytes. **A)** Monocytes were isolated with CD14 negative selection and thereafter stained with a fluorescently labeled CD14 antibody before being analyzed with flow cytometry. The histogram shows one representative experiment out of three. **B).** WKYMVM***-***induced O_2_^−^ production is inhibited by P4Pal_10_ in monocytes. Monocytes purified by CD14 negative selection (10^5^) were incubated (5 min, 37°C) without (solid line) or with P4Pal_10_ (1 µM, PAR4 pepducin, dotted line) before stimulation with WKYMVM (100 nM) as indicated by the arrow. The O_2_^−^ production was measured by chemiluminescence and followed over time. A representative experiment is shown, n = 3. Abscissa: time (Min), Ordinate: chemiluminescence (arbitrary units, MCPM) **C).** Inhibition of WKYMVM-induced O_2_^−^ production in monocytes by the P4Pal_10_ (2 µM) pepducin. The results are presented as remaining activity of WKYMVM (100 nM) or fMLF (100 nM) in the presence of the P4Pal_10_ pepducin, expressed as a percentage of the total production (area under curve (AUC)) induced in the absence of P4Pal_10_ (means ± sd; n = 3). \**p* <0.05. **D).** Neutrophils (10^6^ cells/ml) were incubated with P4Pal_10_ (2 µM, black bar) on ice for 10 min. After which the fluorescently labeled FPR2 agonist Cy5-WKYMVM (1 nM) was added, and incubation was continued for 60 min. Cells pretreated with unlabeled WKYMVM (100 nM; white bar) or without agonist/antagonist (control, total binding) before the addition of Cy5-WKYMVM (1 nM) were included. Data are expressed as percentage of total binding (mean fluorescence intensity ± sd, n = 4).

### The allosteric FFA2R modulator Cmp58 turns P4Pal_10_ into a potent agonist that activates the neutrophil NADPH oxidase

In addition to FPR2, neutrophils express also pattern recognition FFA2R that sense short free fatty acids (22). In contrast to the FPR2 agonists used above, the FFA2R agonist propionate cannot alone trigger an activation of the NADPH-oxidase, but the allosteric FFA2R modulator Cmp58 turns propionate into a potent activator of the NADPH-oxidase (27, 31). Our results not only confirmed the effect of Cmp58 on propionate but also revealed that this allosteric modulator turned also the P4Pal_10_ pepducin into a potent neutrophil activator (Fig 5A). In the presence of Cmp58, the P4Pal_10_-induced O_2_^−^ release was of the same magnitude as that induced by propionate (Fig 5A inset). The P4Pal_10_-induced response was inhibited by an FFA2R antagonist (Fig 5A), clearly showing that the response is mediated through FFA2R. To further characterize the P4Pal_10_-induced activation of the neutrophil NADPH-oxidase, we determined the priming effect of TNFα on this response. In agreement with our previous data with propionate as the activating agonist (20), the P4Pal_10_ response in the presence of Cmp58 was greatly augmented in TNFα primed cells (Fig 5B). In addition, we found that the primed P4Pal_10_ response was sensitive to the Gαq inhibitor YM-254890 (Fig 5B and Fig 5C). The amount of O_2_^−^ production was not significant altered when the order of the Cmp58 and P4Pal_10_ was reversed upon stimulation (inset Fig 5D) but the onset of the response was somewhat delayed (Fig 5D). Taken together, these data clearly demonstrate that together with an allosteric FFA2R modulator, P4Pal_10_ acts as a potent FFA2R agonist that activates the neutrophil NADPH oxidase.

**Figure 5.**
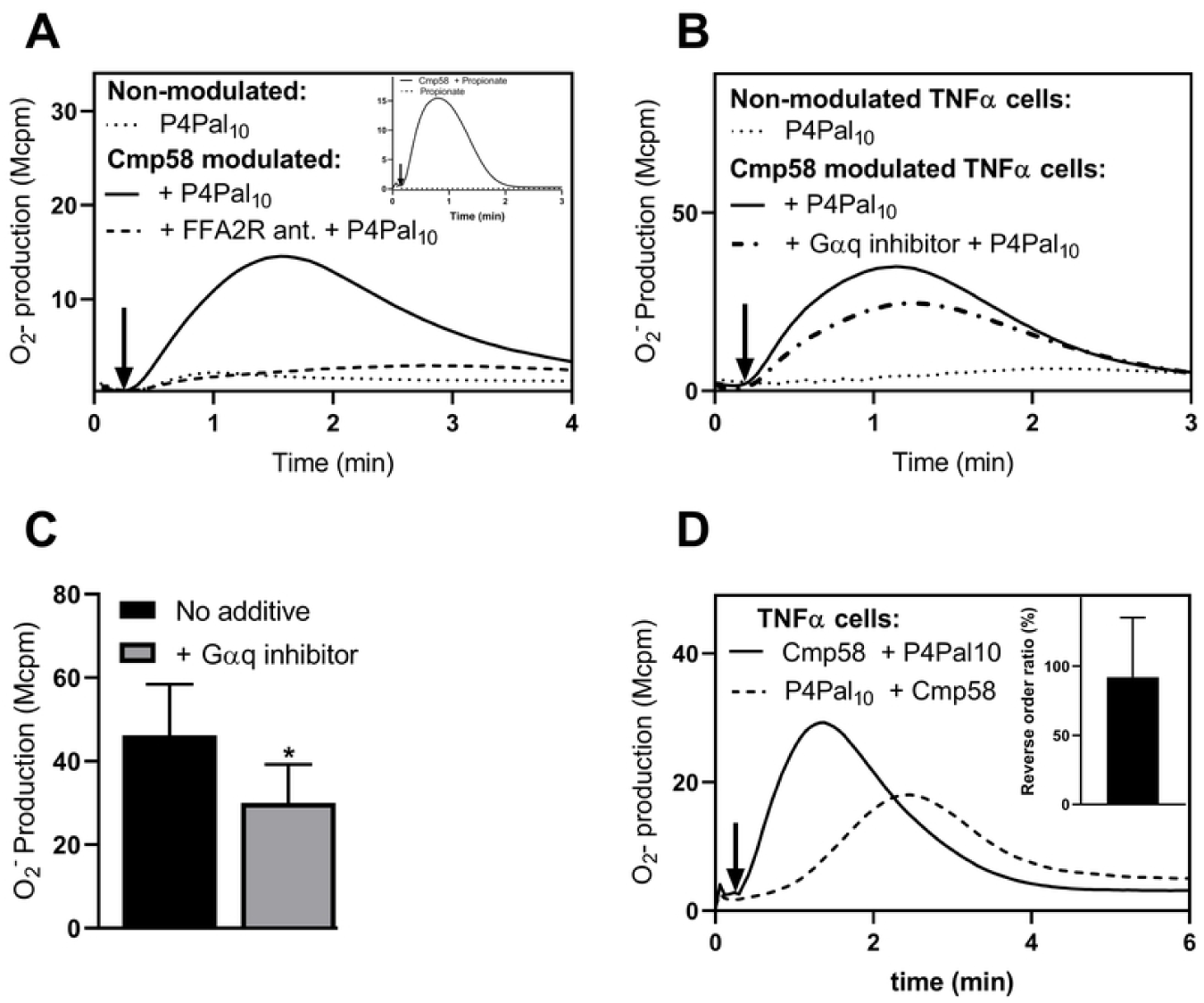
Modulated FFA2R turns P4Pal_10_ into an activator of the NADPH oxidase. **A).** P4Pal_10_ induced O_2_^−^ production in Cmp58 modulated neutrophils Neutrophils (10^5^) were incubated (5 min, 37°C) with or without: Cmp58 (1 µM), P4Pal_10_ (1 µM, PAR4 pepducin) and CATPB (100 nM, FFA2R antagonist) before being stimulated with P4Pal_10_ (1 µM) as indicated in the figure. Inset, Propionate induced O_2_^−^ production in the presence of Cmp58. Neutrophils (10^5^) were incubated (5 min, 37°C) with or without: Cmp58 (1 µM) before being stimulated with propionate (25 µM). The O_2_^−^ production was measured by chemiluminescence and followed over time. A representative experiment is shown, n = 3. Abscissa: time (Min), Ordinate: chemiluminescence (arbitrary units, Mcpm). **B)** P4Pal_10_ induced O_2_^−^ production in TNFα treated and Cmp58 modulated neutrophils Neutrophils (10^5^ pretreated with TNFα (20 ng/mL, 20 min, 37°C) were incubated (5 min, 37°C) with or without: Cmp58 (1 µM), P4Pal_10_ (1 µM, PAR4 pepducin) and YM-254890 (200 nM, Gαq inhibitor) before being stimulated with P4Pal_10_ (1 µM) as indicated in the figure. The O_2_^−^ production was measured by chemiluminescence and followed over time. A representative experiment is shown, n = 3. Abscissa: time (Min), Ordinate: chemiluminescence (arbitrary units, Mcpm). **C).** Inhibition of P4Pal_10_ induced O_2_^−^ production in neutrophils by the Gαq inhibitor YM-254890. The results are presented as activity of P4Pal_10_ (1 µM) with YM-254890 (200 nM, grey bar) or without (black bar) in the presence of Cmp58 (1 µM), expressed as a peak O_2_^−^ production (means ± sd; n = 3). \**p* < 0.05. **D.** Reversed order induced O_2_^−^ production Neutrophils (10^5^ pretreated with TNFα (20 ng/mL, 20 min, 37°C) were incubated (5 min, 37°C) with Cmp58 (1 µM) before being stimulated with P4Pal_10_ (1 µM, solid line) or incubated (5 min, 37°C) with P4Pal_10_ (1 µM, PAR4 pepducin) before being stimulated with Cmp58 (1 µM, dotted line) as indicated in the figure. The O_2_^−^ production was measured by chemiluminescence and followed over time. A representative experiment is shown, n > 3. Abscissa: time (Min), Ordinate: chemiluminescence (arbitrary units, Mcpm). Inset: the neutrophil response induced by Cmp58 and P4Pal_10_ (1 µM respectively) compared to the response induced by the same ligands added in the reversed order and expressed as the ratio of the total production (AUC) values induced by (P4Pal_10_ + Cmp58) / (Cmp58 + P4Pal_10_) (means ± sd, n = 5).

## Discussion

In this study, we have investigated the effects of the presumed Gαq-inhibitory pepducin P4Pal_10_ using human neutrophils and well-characterized agonists specific for Gαq- and Gαi-coupled receptors. Our results show that P4Pal_10_ has no inhibiting effect on signaling transduced by the Gαq-coupled receptors studied, but the pepducin inhibits selectively activation of the Gαi-coupled FPR2 both in human neutrophils and monocytes. In addition, the allosteric FFA2R modulator Cmp58 turns P4Pal_10_ into a potent neutrophil activator and this effect is reciprocal, meaning that the pepducin turns the allosteric modulator into a potent activating agonist. Our findings thus call for further investigation of the precise mechanism of action of pepducins on GPCRs and when it comes to the therapeutic potential for P4Pal_10_ (as well as other pepducins) in different types of diseases, there is a need for a careful evaluation of receptor selectivity. Despite this, both activating and inhibiting lipopeptides including P4Pal_10_ clearly serve as unique tool compounds for further mechanistic studies of GPCR signaling and modulation. GPCR pepducins were introduced some 15 years ago as allosteric receptor modulators with a unique mechanism of action directly involving the cytosolic signaling parts of the receptor (9). According to the suggested model, the peptide part of a pepducin determines the receptor specificity that is fine-tuned by the identity at the level of the amino acid sequence in the peptide part of pepducin and one of cytosolic receptor domains (9), and with an absolute fit between these, the pepducin either activates or inhibits receptor function. The P4Pal_10_ pepducin operates, however, not only on the cognate receptor (PAR4) from which it was derived, but also on receptors that do not share a strong sequence homology with the pepducin (11). The background to the receptor promiscuity of the inhibition mediated by the P4Pal_10_ pepducin was suggested to be linked to the recruitment of the signaling G-protein down-stream of the targeted receptors, with the common denominator being that the sensitive receptors signal through a Gαq containing G-protein (11). Our results in this study clearly show that this mode of action does not apply to human neutrophils as P4Pal_10_ inhibited signaling transduced by the Gαi-coupled FPR2 but not by the Gαq-coupled receptors investigated. In addition, we expanded the P4Pal_10_-targeting GPCRs by including also FFA2R, a receptor designed to recognize short chain fatty acids. The molecular background to the effects of P4Pal_10_ on FPR2 and FFA2R cannot be fitted into the original pepducin concept stating that the peptide sequence of a receptor modulating pepducin, should be present also in one of the cytosolic parts of the targeted receptor. There are no sequence similarities between P4Pal_10_ (peptide sequence (SGRRYGHALR)) and any of the cytosolic parts of FPR2 and FFA2R. The basis for our study was that P4Pal_10_ was claimed to be an inhibitor of Gαq signaling, with an inhibition profile across multiple receptor subtypes, of which some have no or very limited sequence similarity in their intracellular domains, with the third intracellular loop of PAR4 from which the amino acid sequence in the P4Pal_10_ pepducin was derived (32). The precise mechanism by which P4Pal_10_ affects the diverse set of Gα_q_-coupled receptors (11), thus, remains elusive, but according to the pepducin concept, P4Pal_10_ should affect some unknown but critical intra-receptor or receptor–G protein interaction site that is necessary for the down-stream signaling promoted selectively by a Gαq protein. It is clear from the data presented, that the putative Gαq inhibitory pepducin P4Pal_10_ does not inhibit the Gαq-coupled neutrophil receptors PAFR and P2Y_2_R in human neutrophils, suggesting that the unknown critical interaction site required for inhibition is missing in these receptors.

More importantly, we present some novel modes of action of P4Pal_10_; it antagonizes the Gαi-coupled FPR2 and activates the allosterically modulated FFA2Rs. It should also be noticed, that the effect of Cmp58 on the P2Pal_10_ induced response was the same as when the order of addition was reversed and the pepducin was used as sensitizing ligand. This type of allosteric interaction is reciprocal in nature and the results presented are, thus, in agreement with the reciprocity characterizing receptor allosterism. As mentioned, the pepducin dogma states that active pepducins are expected to allosterically modulate receptor function through an interaction with an intracellular receptor-specific binding site; the fatty acid anchors the pepducin to the cell membrane and allow the peptide part to flip over and enter the cytosol to interact and activate or inhibit signaling by its cognate receptor (9). We have in a series of earlier studies presented data that strongly challenge the dogma when showing that, i) even though and FPR2 derived pepducin is a highly selective activating FPR2 agonist, its activity is inhibited by conventional orthosteric antagonists, ii) there is no direct link between the amino acid sequence in the targeted receptor and in the activating/inhibiting pepducin and, iii) pepducins with peptide sequences derived from intracellular domains present in several GPCRs hijack FPR2 (33-35). Irrespective of the precise mode of action of P4Pal_10_, we now add the putative Gαq inhibitor P4Pal_10_ to the list of FPR2 hijacking pepducins, and this together with earlier published data raise questions not only about the general mode of action of pepducins but also about FPR2 and FFA2R as pattern recognition receptors for some type of endogenous or microbe-derived, lipid-substituted peptides that might represent an additional molecular pattern that is recognized by and activate or inhibit FPRs/FFARs.

In summary, we show that the putative Gαq inhibitory pepducin P4Pal_10_ does not inhibit the Gαq-coupled PAFR and P2Y_2_R expressed in in neutrophils but the lipopeptide selectively antagonizes the Gαi-coupled FPR2. In addition, we show that an allosteric FFA2R modulator turns P4Pal_10_ into an activator of the O_2_^−^ generating neutrophil NADPH oxidase.

## Author contributions

AH and SL collected and analyzed data. CD and HF supervised the study. All authors discussed the results and wrote the paper together.

## Acknowledgements

This work was supported by the Swedish Research Council, King Gustaf V 80-Year Foundation, the Swedish government under the ALF-agreement, the Clas Groschinsky Memorial Foundation, and the Ingabritt and Arne Lundberg Foundation.

